# Distinct mechanisms of plant immune resilience revealed by natural variation in warm temperature-modulated disease resistance among *Arabidopsis* accessions

**DOI:** 10.1101/2023.11.15.565111

**Authors:** Christina A. M. Rossi, Dhrashti N. Patel, Christian Danve M. Castroverde

## Abstract

Elevated temperature suppresses production of the key plant defence hormone salicylic acid (SA). Heat-mediated SA suppression and resulting plant vulnerability are due to downregulated expression of *CALMODULIN BINDING PROTEIN 60-LIKE G* (*CBP60g*) and *SYSTEMIC ACQUIRED RESISTANCE DEFICIENT 1* (*SARD1*), which encode master regulators of plant immunity. However, previous studies in *Arabidopsis thaliana* plants have primarily focused on the accession Columbia-0 (Col-0), while the mechanisms governing the intraspecific variation in *Arabidopsis* immunity under elevated temperature have remained unknown. Here we show that BASIC HELIX LOOP HELIX 059 (bHLH059), a thermosensitive SA regulator at non-stress temperatures, does not regulate immune suppression under warmer temperatures. In agreement, temperature-resilient and -sensitive *Arabidopsis* accessions based on disease resistance to the bacterial pathogen *Pseudomonas syringae* pv. *tomato* (*Pst*) DC3000 did not correlate with *bHLH059* sequence polymorphisms. Instead, we found that different temperature-resilient accessions exhibit varying *CBP60g* and *SARD1* expression profiles, potentially revealing both *CBP60g/SARD1*-dependent and independent mechanisms of plant immune resilience to warming temperature. Collectively, this study has unveiled the intraspecific diversity of *Arabidopsis* immune responses under warm temperatures. Our dissection of mechanisms underlying temperature-modulated plant immunity could aid in predicting plant responses to climate change and provide foundational knowledge for climate-resilient crop engineering.

## Introduction

Plant health is critical to maintaining global food and bioenergy security; however, plant disease outbreaks are only becoming more severe and prevalent with climate change (Savary et al., 2019; Deutsch et al., 2018; Chaloner et al., 2021). The relationship between host plants and disease-causing pathogens with dynamically changing environmental conditions is well-synthesized by the “plant disease triangle” paradigm (Stevens, 1960; Francl, 2001). For example, plants experiencing sub-optimal environmental conditions, such as heat or water stress, exhibit weaker immune responses and therefore become more susceptible to pathogen infection (Velásquez et al., 2018; Cohen and Leach, 2020; Desaint et al., 2021; Singh et al., 2023; Roussin-Leveillee et al., 2023).

The cross-kingdom interaction between *Arabidopsis thaliana* plants and the bacterial pathogen *Pseudomonas syringae* pv. *tomato* (*Pst*) DC3000 has been used extensively as a model system to study host-pathogen interactions and the plant disease triangle (Ausubel et al., 1995; Xin and He, 2013; Velásquez et al., 2018). As shown in *Arabidopsis* and various other plant species, plants elicit a two-pronged and interlinked immune response during pathogen attack: (1) pathogen-associated molecular pattern (PAMP)-triggered immunity (PTI) and (2) effector-triggered immunity (ETI) (Bigeard et al., 2015; Zhou and Zhang et al., 2020; Yuan et al., 2021; Ngou et al., 2022). A key plant defence hormone mediating convergent immune signalling is salicylic acid (SA), which is crucial for local and systemic defences against biotrophic and hemibiotrophic pathogens (Zhang and Li, 2019; Peng et al., 2021; Rossi et al., 2023). During pathogen challenge, endogenous SA production in plants is important in amplifying both PTI and ETI (Tateda et al., 2014; Liu et al., 2020; Yang et al., 2021; Shields et al., 2022).

Previous studies have demonstrated that elevated temperatures inhibit specific components of the plant immune system, including PTI, ETI, and SA production (De Jong et al., 2002; Wang et al., 2009; Zhu et al., 2010; Cheng et al., 2013; Menna et al., 2015; Huot et al., 2017; Velasquez et al., 2018; Kim et al., 2021; Kim et al., 2022; Shields et al., 2023). In particular, ISOCHORISMATE SYNTHASE 1(ICS1)-mediated SA biosynthesis is downregulated at elevated temperature (Malamy et al., 1992; Huot et al., 2017; Castroverde and Dina, 2021; Shields et al., 2023). Heat suppression of the SA pathway is governed through the thermosensitive, rate-limiting transcription of the master immune regulatory genes *CBP60g* and *SARD1* (Kim et al., 2022). Recently, a study uncovered that warmer temperatures (28°C) reduce the formation of intranuclear GUANYLATE BINDING PROTEIN-LIKE 3 (GBPL3) defence-activated biomolecular condensates (GDACs), thereby abolishing the recruitment of the general transcriptional machinery to the *CBP60g* and *SARD1* genetic loci (Kim et al., 2022). Overall, temperature has a profound impact on the plant immune system through various levels of genetic regulation.

In addition to changing environmental conditions, SA production is also influenced by the profound intraspecific diversity of the *Arabidopsis* pangenome (van Leeuween et al., 2007; Bechtold et al., 2010; Yang et al., 2015; Velasquez et al., 2017; Bruessow et al., 2021). At non-stress temperatures (16°C-22°C), Bruessow *et al*. (2021) discovered remarkable natural variation in basal (uninduced) SA accumulation among numerous *Arabidopsis* accessions. Through a genome-wide association study (GWAS) of >100 natural *A. thaliana* accessions, thermoresponsive basal SA production was found to be mediated by the transcription factor BASIC HELIX-LOOP-HELIX 059 (bHLH059; Bruessow et al., 2021). Bruessow *et al*. (2021) found two significant SNPs that distinguish *Arabidopsis* accessions into thermosensitive or thermoresilient phenogroups based on SA levels. Furthermore, the authors also identified that *bHLH059* transcripts increased at 22°C compared to 16°C in temperature-sensitive accessions (e.g., Col-0); however, *bHLH059* gene expression remained unchanged in temperature-resilient accessions (Bruessow et al., 2021).

A critical gap in the current literature is that the *Arabidopsis* immune variation at elevated temperature and its underlying mechanisms have not been fully dissected. Here we show extensive intraspecific diversity among *Arabidopsis* accessions in terms of disease susceptibility to the bacterial pathogen *Pst* DC3000 between ambient (23°C) and moderately elevated temperatures (28°C). Unlike at the non-stress temperature range, we further demonstrate that this natural variation is independent of bHLH059 but exhibits accession-specific dependence on the expression of the immune regulatory genes *CBP60g* and *SARD1*. Collectively, leveraging the naturally occurring variation within the *A. thaliana* species may prove to be an effective strategy to better understand how plant immune systems respond to our changing climate.

## Materials and Methods

### Plant materials and growth conditions

*Arabidopsis* mutants and accessions used in this study were obtained from the *Arabidopsis* Biological Resource Centre, The Ohio State University (see Supplementary Table 1). Seeds were surface-sterilized with 70% ethanol, rinsed three times with autoclaved water and then stratified in 0.1% agarose at 4°C for 3 days to promote uniform germination (Rivero et al., 2013). Seeds were sown onto autoclaved soil comprised of equal parts ProMix PGX or BX (Plant Products, Ancaster, Ontario), Turface (Turface Athletics, Buffalo Grove, IL), and Vermiculite (Therm-O-Rock East, New Eagle, PA), supplemented with MiracleGro (The Scotts Company, Mississauga, ON). Plants were incubated in environmentally controlled growth chambers with the following conditions: 23°C temperature, 60% relative humidity, 12h day/12h night cycle, and 80-100 μmol m^-2^ s^-1^ light conditions.

### Plant infection and disease assays

Three leaves of 4-to 5-week-old *Arabidopsis* plants were syringe-infiltrated with the *Pst* DC3000 suspension (OD600=0.001), which was prepared as previously described (Huot et al., 2017; Kim et al., 2022; Shields et al., 2023). Plants were then incubated at 23°C (normal) or 28°C (elevated) with the same relative humidity and light cycling conditions. At 3 days post-inoculation (dpi), in planta bacterial quantification was performed as described previously (Huot et al., 2017). Briefly, circular leaf punches were collected from the infiltrated leaves and homogenized in 0.25 mM MgCl_2_ with a TissueLyser II (Qiagen, Toronto, ON). Samples were then serially plated onto LM media containing rifampicin and incubated at 21-24°C overnight. Colony-forming units (CFUs) were counted two days later, and bacterial levels were reported as log CFU/cm^2^.

### Gene expression analyses

Leaves of 4-to 5-week-old *Arabidopsis* plants were syringe-infiltrated with mock solution (0.25 mM MgCl_2_) or *Pst* DC3000 suspension (OD600=0.001) as described above. Plants were then incubated at 23°C (normal) or 28°C (elevated) with the same relative humidity and light cycling conditions. Gene expression levels were quantified based on a previously published protocol (Kim et al., 2022) with slight modifications. At 1 dpi, total RNA was extracted from flash-frozen plant tissues using the Qiagen Plant RNeasy Mini Kit (Qiagen, Toronto, ON) and cDNA was synthesized using qScript cDNA super mix (Quantabio, Beverly, MA) based on manufacturers’ recommendations. Real-time quantitative polymerase chain reaction (qPCR) was performed using PowerTrack SYBR Green master mix (Applied Biosystems, Waltham, MA) or ChamQ Universal SYBR qPCR Master Mix (Vazyme Biotech, City of Dover, DE). qPCR amplification was performed using the Applied Biosystems QuantStudio3 platform (Life Technologies), and individual Ct values were determined for target genes (*CBP60g, SARD1*) and the internal control gene (*PP2AA3*) (Huot et al., 2017; Kim et al., 2022). Gene expression values were reported as 2^−ΔCt^, where ΔCt is Ct_target gene_–Ct_*PP2AA3*_. qPCR was carried out with three technical replicates for each biological sample. Primers used for qPCR are shown in Supplementary Table 2.

### Statistical analyses

Statistical analyses of bacterial quantification and gene expression values were carried out using R (R Core Team, 2023) or Prism (GraphPad). Specific statistical tests for each experiment are detailed in the respective figure captions.

## Results

### The transcription factor bHLH059 is not involved in temperature-sensitive *Arabidopsis* immunity under warm temperatures

Because bHLH059 acts as a thermoresponsive SA regulator at non-stress temperatures between 16°C and 22°C (Bruessow et al., 2021), we tested whether bHLH059 also controls the natural variation of temperature-modulated immunity at warm temperatures in the 23°-28°C range. As shown in Figure 1A-B, the reference accession Col-0 expectedly exhibited temperature-sensitive disease resistance in terms of pathogen levels and disease symptoms, consistent with previous studies (Wang et al., 2009; Mang et al., 2012; Huot et al., 2017; Kim et al., 2022). Both *blhh059* mutant alleles similarly showed thermosensitive disease susceptibility to *Pst* DC3000, with higher bacterial levels and disease symptoms at 28°C than at 23°C. The two *bhlh059* mutants were also evaluated for gene expression levels of *CBP60g* and *SARD1*, which were recently reported to be the rate-limiting nodes in warm temperature suppression of the plant immune system (Kim et al., 2022). As shown in Figure 1C-D, Col-0 showed temperature-sensitive *CBP60g* and *SARD1* gene expression as expected (Kim et al., 2022). Similar to Col-0, both *bhlh059* mutants also showed temperature-sensitive *CBP60g* and *SARD1*gene expression, with lower transcript levels at 28°C compared to 23°C after pathogen infection. Taken together, these results indicate that the transcription factor bHLH059 does not play an essential role in controlling warm temperature-modulated immunity in *Arabidopsis* plants.

**Fig. 1.**
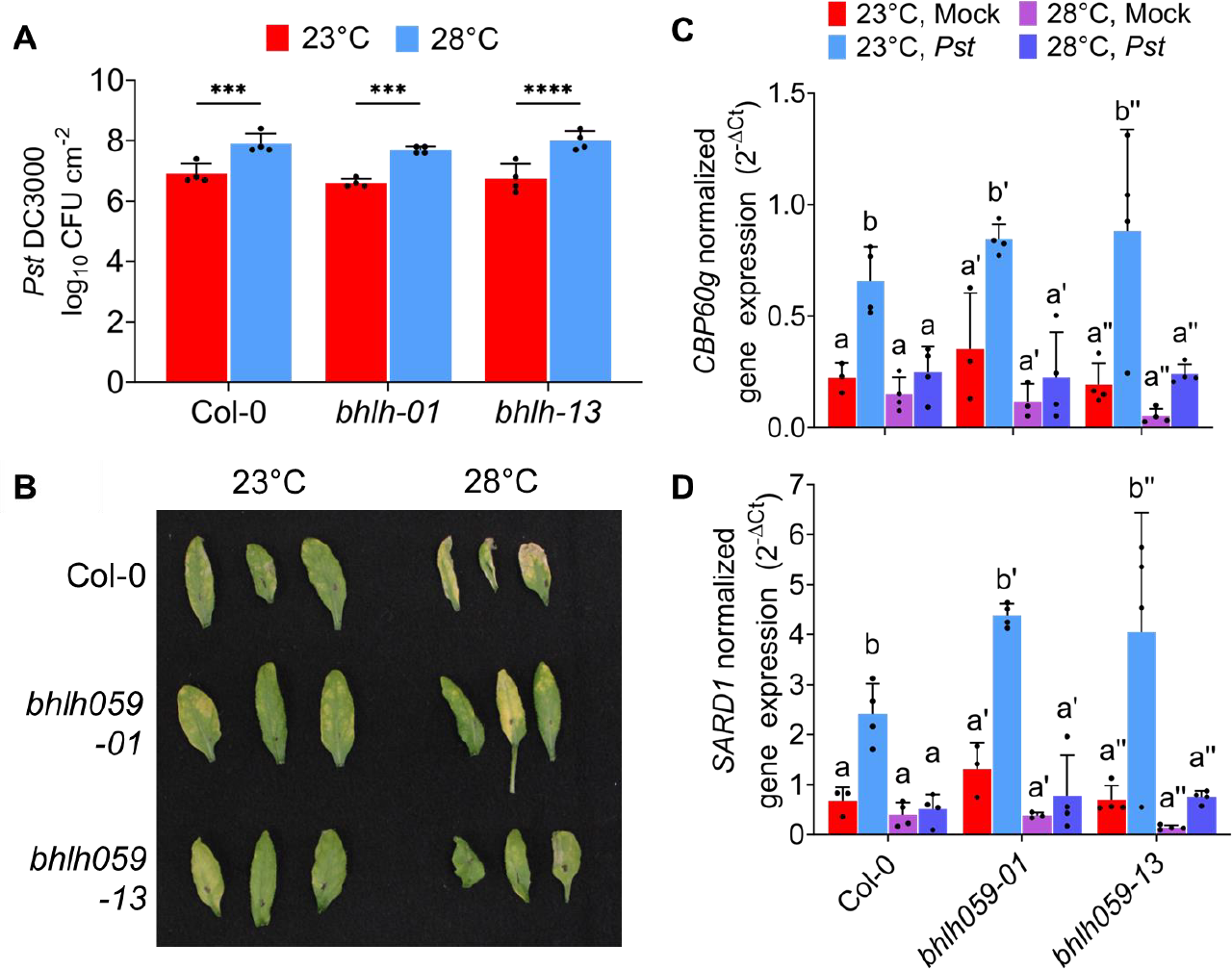
Disease susceptibility phenotypes and defence gene expression profiles of the *Arabidopsis bhlh059* mutants. Leaves of four-week-old *bhlh059* mutants of *A. thaliana* were infected with *Pst* DC3000 (OD_600_ = 0.001) and incubated at 23°C or 28°C for three days after inoculation. (A) In planta bacterial levels were quantified at 3 days post-inoculation (dpi). Results shown are the Log_10_(CFU)cm^-2^ means ± SD (n = 4 individual plants from one representative experiment). (B) Three leaves from one representative plant were harvested at 3 dpi and photographs were taken to show disease symptoms. (C-D) At 1 dpi, plant leaf total RNA was extracted and *CBP60g* or *SARD1* gene expression levels (normalized to *PP2AA3* internal control) were determined. Results shown are the means ± SD (n =3 or 4 from one representative independent experiment). Data were analyzed with two-way ANOVA and Tukey’s multiple comparisons test (*p* < 0.05). Statistically significant values are indicated by asterisks (*p* < 0.05, ** *p* < 0.1, *** *p* < 0.001, **** *p* <0.0001) or different letters. This experiment was repeated at least two times with reproducible results.

### Impact of elevated temperature on disease resistance in various natural accessions of *A. thaliana*

As shown above and in previous studies, the reference accession *A. thaliana* Col-0 exhibits temperature-sensitive immune responses at non-stress temperatures (Li et al., 2020; Bruessow et al., 2021) and at warm temperatures (Huot et al., 2017; Kim et al., 2022). To investigate the intra-species diversity of *Arabidopsis* immunity at elevated temperature, disease resistance assays were performed by inoculating four-week-old *A. thaliana* natural accessions (including Col-0) with virulent *Pst* DC3000. As shown in Figure 2A, certain accessions (Col-0, Ler, Fei-0, and Est-1) had significantly higher bacterial levels at 28°C than at 23°C (i.e. temperature-sensitive), while other accessions (Cvi-0, NW-1, Se-0, Ei-2, NFA-8, Sf-2, Bay-0, and Mz-0) did not have significantly different *Pst* DC3000 pathogen levels between the two temperatures (i.e. temperature-resilient). It is important to note that there was considerable natural variation in basal disease resistance within each group. Among the temperature-sensitive accessions, Col-0 had the overall highest bacterial levels, while Est-1 had the lowest bacterial levels (Fig. 2A). On the other hand, among the temperature-resilient accessions, Cvi-0 had the highest overall *Pst* DC3000 levels, while Mz-0 had the lowest (Fig. 2A). Similar to the in planta bacterial levels, there was also considerable variation in disease symptom development among these accessions as shown in Fig. 2B. Overall, these results indicate that elevated temperature differentially impacts the defence responses of various *A. thaliana* accessions to *Pst* DC3000.

**Fig. 2.**
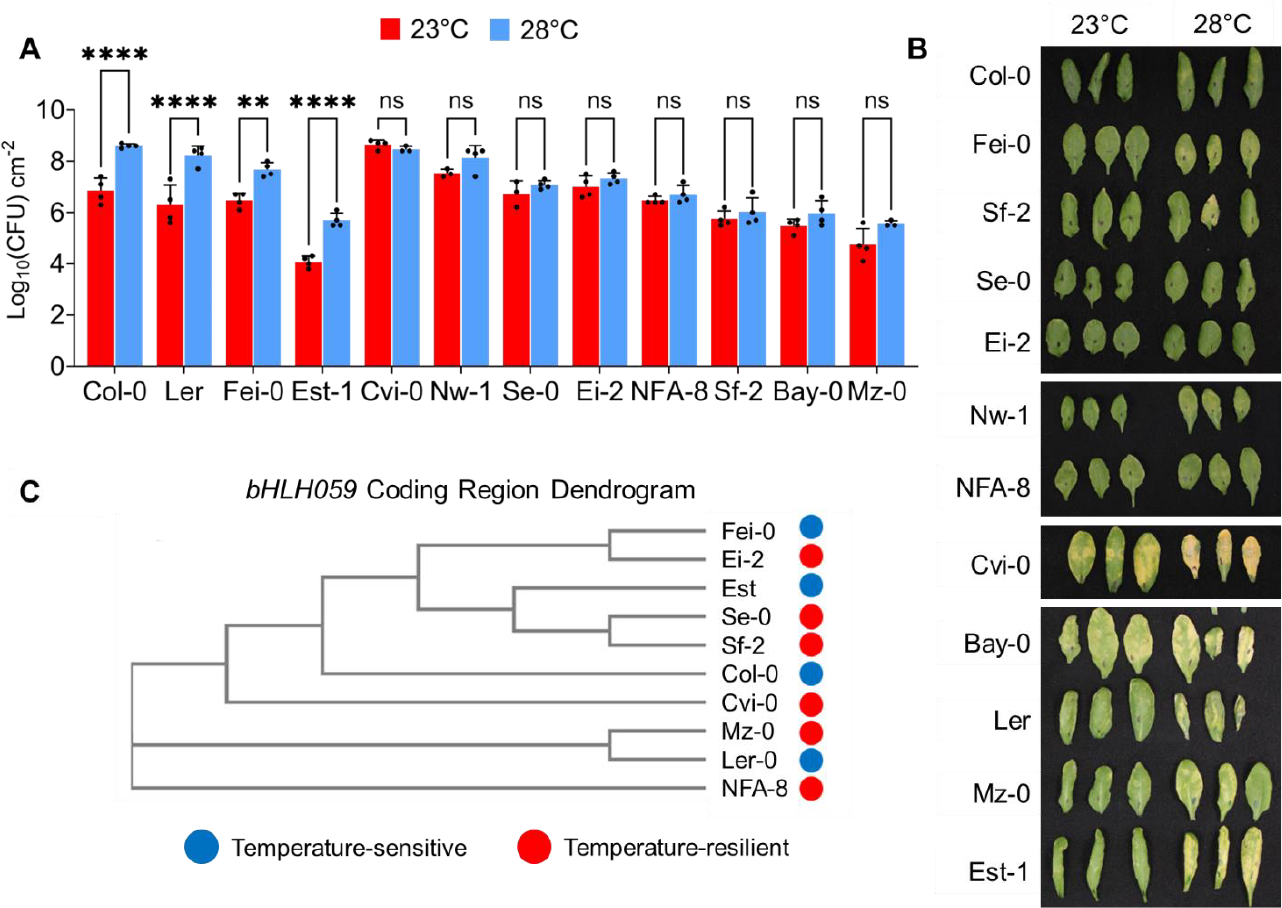
Natural variation in thermoregulated disease resistance to *Pst* DC3000 and *bHLH059\* sequence diversity in different natural accessions of *Arabidopsis thaliana*. (A) Leaves of four-week-old accessions of *A. thaliana* were infected with *Pst* DC3000 (OD_600_ = 0.001) and incubated at 23°C or 28°C for three days after inoculation. In planta bacterial levels were quantified at 3 days post-inoculation (dpi). Results shown are the Log_10_(CFU)cm^-2^ means ± SD (n = 3 or 4 individual plants from one representative experiment). Data were analyzed with one-way ANOVA and Tukey’s multiple comparisons test (*p* < 0.05). Statistically significant values are indicated by asterisks (*p* < 0.05, ** *p* < 0.1, *** *p* < 0.001, **** *p* <0.0001). The experiment was independently replicated three times with reproducible results. (B) Three leaves from one representative plant were harvested at 3 dpi and photographs were taken to show disease symptoms. (C) Known *bHLH059* DNA coding regions of ten natural Arabidopsis accessions were clustered in Clustal Omega into a dendrogram as a neighbour-joining tree without distance corrections.

Under the non-stress temperature range from 16°-22°C, distinct polymorphisms in the SA regulator bHLH059 controls natural variation in temperature-modulated basal SA levels (Bruessow et al., 2021). To better understand if the intraspecific variation under the warmer temperature (23°-28°C) also correlates with bHLH059 sequence polymorphisms, we clustered different accessions based on the *bHLH059* coding region. As shown in Figure 2C, hierarchical clustering of *bHLH059* protein-coding DNA sequences did not strictly separate the different accessions based on their disease resistance phenotypes determined in Figure 1A-B. This suggests that bHLH059-independent mechanisms govern differential immune resilience or sensitivity to elevated temperatures in *Arabidopsis* plants.

### Differential *CBP60g* and *SARD1* gene expression in *Arabidopsis* accessions with temperature-resilient disease resistance

To further investigate the underlying mechanisms of thermoregulated plant immune variation, we examined gene expression levels of two master immune regulatory genes *CBP60g* and *SARD1* (Wang et al., 2009; Zhang et al., 2010; Wang et al., 2011) in three representative temperature-resilient accessions (Se-0, Sf-2, NFA-8) and reference accession Col-0. This aimed to establish if these two genes are the driving force behind temperature-resilient SA levels and immunity in naturally occurring *Arabidopsis* accessions. As shown in Figure 3A-B, Col-0 expectedly showed temperature-sensitive *CBP60g* and *SARD1* gene expression, with significantly lower transcript levels at 28°C compared to 23°C after pathogen infection (Kim et al., 2022; Shields et al., 2023). Interestingly, the temperature-resilient accessions Se-0 and NFA-8 behaved similarly as Col-0 in terms of defence gene expression, showing significantly lower levels of pathogen-induced *CBP60g* and *SARD1* at 28°C compared to 23°C (Figure 3A-B). In contrast, the temperature-resilient accession Sf-2 showed temperature-resilient *CBP60g* and *SARD1* gene expression profiles, with similar pathogen induction of transcript levels between 23°C and 28°C (Fig. 3A-B). To clarify whether genetic polymorphisms in these master immune regulators also contribute to temperature-resilient immunity, we clustered the *CBP60g* and *SARD1* coding sequences of the full accession panel in Figure 3C and D. However, there was no association between coding region sequence polymorphisms in the different accessions and their level of disease thermosensitivity. Overall, these results suggest that there may be both *CBP60g/SARD1* expression-dependent (as shown in Sf-2) and -independent (as shown in Se-0 and NFA-8) mechanisms conferring immune resilience across the *Arabidopsis* pangenome.

**Fig. 3.**
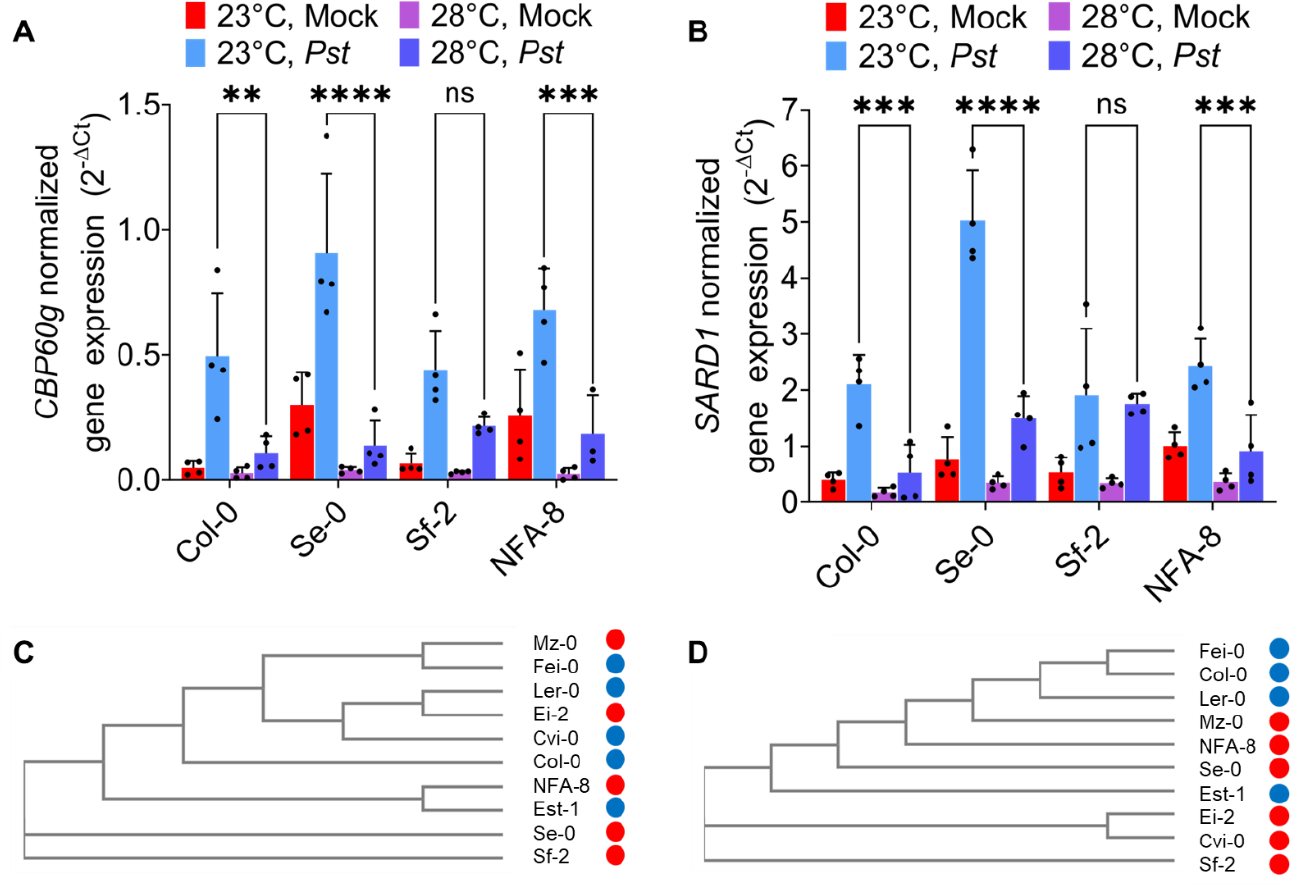
Variation in *CBP60g* and *SARD1* gene expression profiles and sequence polymorphisms in representative temperature-resilient Arabidopsis accessions. (A-B) Four-week-old *Arabidopsis* plant leaves were infiltrated with 0.25mM MgCl_2_ mock solution or *Pst* DC3000 suspension (OD_600_ = 0.001). Plants were then incubated at either 23°C or 28°C. One day post-infection (1 dpi), plant tissues were collected and *CBP60g* (A) or *SARD1* (B) gene expression levels (normalized to *PP2AA3* internal control) were determined. Results shown are the means ± SD (n= 4 from one representative independent experiment). Data were analyzed with two-way ANOVA and Tukey’s multiple comparisons test (*p* < 0.05). Statistically significant values are indicated by asterisks (* *p* < 0.05, ** *p* < 0.1, *** *p* < 0.001, **** *p* <0.0001). (C-D) DNA coding region clustering of natural *Arabidopsis* accessions were conducted in Clustal Omega as a neighbour-joining tree without distance corrections for both *CBP60g* (C) and *SARD1* (D). Red represents a temperature-resilient accession, while blue represents a temperature-sensitive accession.

## Discussion

Rising global temperatures associated with climate change have a profound impact on plant health, including effective immune responses to infectious diseases (Savary et al., 2019; Deutsch et al., 2018; Velásquez et al., 2018; Chaloner et al., 2021; Singh et al., 2023). However, the extent of intraspecific variation in plant immunity at warm temperatures has remained unexplored. In this study, we demonstrated extensive variation in temperature-modulated disease susceptibility among natural *A. thaliana* accessions at elevated temperatures (23°-28°C). In striking contrast to the non-stress temperature range (16°-22°C), this natural variation is independent of the recently identified SA regulator bHLH059 (Bruessow et al., 2021). Instead, temperature-resilient *Arabidopsis* immunity can be mediated by distinct regulatory mechanisms that are either dependent or independent on sustained expression of *CBP60g* and *SARD1*, which encode master transcription factors of SA biosynthesis (Wang et al., 2009; Zhang et al., 2010; Wang et al., 2011; Sun et al., 2015; Kim et al., 2022).

Using a panel of *Arabidopsis* accessions based on previous studies (Bruessow et al., 2021; Sanchez-Bermejo et al., 2015), disease resistance assays to the bacterial pathogen *Pst* DC3000 revealed two key groups: (1) temperature-sensitive accessions, with greater bacterial growth *in planta* at 28°C compared to 23°C, and (2) temperature-resilient accessions, with similar levels of *in planta* bacterial growth at 28°C and 23°C. Temperature-sensitive accessions included Col-0, Ler, Fei-0, and Est-1, while temperature-resilient accessions included Se-0, Ei-2, NFA-8, Sf-2, Cvi-0, Bay-0, and Mz-0. A previous study has found that certain accessions experience temperature-sensitive or -resilient immune responses at non-stress temperatures between 16°C and 22°C (Bruessow et al., 2021). However, Bruessow *et al*. (2021) did not evaluate how these various accessions respond to warmer temperatures (e.g., 28°C), which had remained a major knowledge gap until our study. Comparing the non-stress (16°-22°C) and elevated temperature ranges (23°-28°C), certain differences in thermoregulated immunity are apparent. For example, while Bruessow et al. (2021) found that Bay-0 exhibited thermosensitive disease resistance between 16°C and 22°C, we found that this accession is thermoresilient at warmer temperatures. In contrast, Ler was determined to have better immunity at 22°C compared to 16°C (Bruessow et al., 2021), but this accession had higher disease vulnerability at 28°C than at 23°C as shown in our present study. Collectively, our findings (together with those in previous work) reveal the naturally occurring intra-species diversity of disease phenotypes conferred by the *Arabidopsis* pangenome. Future research could expand this study by examining a broader panel of natural accessions.

Having established the natural variation in temperature-sensitivity or -resilience of *Arabidopsis* disease resistance to the pathogen *Pst* DC3000, we then sought to determine the underlying mechanism regulating this intraspecific diversity. We investigated the role of the transcription factor bHLH059, since it has been implicated in thermoregulation of immunity at non-stress temperatures (16°C vs. 22°C; Bruessow et al., 2021). As shown by Bruessow et al. (2021), *bhlh059* mutants exhibit temperature-insensitive disease resistance and basal SA levels between 16°C and 22°C. Surprisingly, our results showed that *bhlh059* mutants are temperature-sensitive in terms of *Pst* DC3000 disease susceptibility and SA-related defence gene expression in the 23°C to 28°C temperature range like the reference accession Col-0 (Kim et al., 2022). Furthermore, thermoresilient and thermosensitive *Arabidopsis* accessions did not correlate with *bHLH059* sequence polymorphisms. Taken together, these findings underscore that bHLH059 is not involved in thermoregulated immunity under warmer temperatures, suggesting distinct regulatory mechanisms depending on the temperature range.

Because we eliminated bHLH059 as a potential regulator of *Arabidopsis* plant immunity in the elevated temperature range (23°C-28°C), we examined the expression of *CBP60g* and *SARD1* in representative temperature-sensitive (Col-0) and temperature-resilient accessions (Se-0, Sf-2, NFA-8). *CBP60g* and *SARD1* are key thermoresponsive genes that encode master transcription factors that are rate-limiting for immune resilience to warming conditions (Kim et al., 2022). The temperature-resilient accession Sf-2 showed sustained *CBP60g* and *SARD1* at both normal and elevated temperatures, which is consistent with our previous study showing that plants with constitutive *CBP60g* or *SARD1* expression exhibit immune resilience to changing temperatures (Kim et al., 2022). Surprisingly, the other two temperature-resilient accessions Se-0 and NFA-8 showed temperature-sensitive *CBP60g* and *SARD1* levels (i.e., still suppressed at warm temperature). These thermosensitive defence gene expression profiles diverge from their disease susceptibility phenotypes and *in planta* pathogen levels. In addition to defence gene expression profiling, our sequence analyses found no distinguishing nucleotide or amino acid polymorphisms in *CBP60g* and *SARD1* to delineate the naturally temperature-resilient or sensitive immunity in representative *Arabidopsis* accessions. This suggests *CBP60g/SARD1* expression-independent immune resilience in Se-0 and NFA-8, which will be important directions to pursue in future studies. These *CBP60g/SARD1*-independent mechanisms may play a role in providing robust disease resistance of certain *A. thaliana* accessions to *Pst* DC3000. In the future, GWAS could be a powerful tool to identify and characterize novel genetic factors involved in the temperature modulation of plant immunity across the entire *Arabidopsis* pangenomic landscape.

Taken together, our study has started to shed light on current knowledge gaps regarding plant immune variation under climate warming conditions. The identification of *Arabidopsis* accessions with temperature-resilient and -sensitive defences will help further uncover molecular components underpinning temperature-sensitive plant immunity. Our findings could aid in predicting how plants respond to our changing climate, potentially laying the groundwork for engineering disease-resistant and climate-resilient plants, which will have broad impacts on agricultural and natural ecosystems.

## Acknowledgements

We thank Castroverde lab members and Dr. Sheng Yang He (Duke University) for meaningful discussion and feedback. We are also grateful to Max Pottier and Gena Braun for their instrumentation support. C.D.M.C. is supported by research funding from the NSERC Discovery Grant, NSERC Research Tools and Instrumentation, Canada Foundation for Innovation, Ontario Research Fund and Wilfrid Laurier University start-up and equipment funds. C.A.M.R. is supported by the Ontario Graduate Scholarship and NSERC Canada Graduate Scholarship. Our research is located on the shared traditional territory of the Neutral, Anishinaabe, and Haudenosaunee peoples.

## Authors’ contributions

C.D.M.C conceptualized and supervised the study. C.A.M.R. performed most of the experiments. D.N.P. was involved with conducting independent experimental replicates of the gene expression analyses. Everyone analyzed the data. C.A.M.R. and C.D.M.C wrote the paper with input from all authors.

## Notes

### Competing Interest Statement

The authors have declared no competing interest.

