## Supplementary Information for "Distinct mechanisms of plant immune resilience revealed by natural variation in warm temperature-modulated disease resistance among *Arabidopsis* accessions"

**Running Title:** Natural variation of *Arabidopsis* plant immune resilience

**Supplementary Table 1. List of Arabidopsis mutants and accessions.**

| Accession or mutant | Description | Reference |
|---|---|---|
| Col-0 | Wild-type germplasm or reference accession | Huot et al., 2017; Kim et al., 2022 |
| *bhlh-01* | SALK_010825C; CS68557 | Bruessow et al., 2021 |
| *bhlh-13* | SALK_135303; CS67000 | |
| Bay-0 | Selected based on Bruessow *et al.* (2021), as these accessions belong to distinct phenogroups for basal SA levels at optimal temperatures. | |
| Ei-2 | | |
| Se-0 | | |
| Est-1 | | |
| Fei-0 | | |
| Mz-0 | | |
| NFA-8 | | |
| Ler | | |
| Sf-2 | Selected based on ePlant (Waese et al., 2017; http://bar.utoronto.ca/eplant/) transcriptome datasets showing high SA gene expression in these two accessions | This study |
| NW-1 | | |
| Cvi-0 | Exhibits temperature-insensitive growth in contrast to Col-0 (Sanchez-Bermejo et al., 2015), which could potentially correlate with temperature-insensitive SA biosynthesis and immunity. | Sanchez-Bermejo et al., 2015 |

**Supplementary Table 2. List of primers used in this study.**

| Primer Name | Sequence | Reference |
|---|---|---|
| *PP2AA3_forward* | GGTTACAAGACAAGGTTCACTC | Huot et al., 2017 |
| *PP2AA3_reverse* | CATTCAGGACCAAACTCTTCAG | |
| *CBP60g_forward* | TCGTGGACGCCACCACAAACA | Kim et al., 2017 |
| *CBP60g_reverse* | TCAGCGTTCAGCGGCACGAG | |

| | | |
|---|---|---|
| *SARD1_forward* | TCGAGTTGGATTCGTAGCCG | Kim et al., 2017 |
| *SARD1_reverse* | TCGCTTCAGTCATCGCTTCA | |
